# Biomechanical and Compositional Changes in the Murine Uterus with Age

**DOI:** 10.1101/2024.09.28.615592

**Authors:** Mari J.E. Domingo, Triniti N. Vanoven, Raffaella De Vita, Maria E. Florian Rodriguez, Kristin S. Miller, Isaac J. Pence

**Affiliations:** Department of Bioengineering, University of Texas at Dallas, Richardson, TX 75080, USA; Department of Biomedical Engineering, University of Texas Southwestern Medical Center, Dallas, TX, 75390, USA; Department of Mechanical Engineering, Virginia Tech, Blacksburg, 24061, VA, USA; Department of Obstetrics and Gynecology, University of Texas Southwestern Medical Center, Dallas, TX, 75390, USA; Department of Mechanical Engineering, University of Texas at Dallas, Richardson, TX 75080, USA; Charles and Jane Pak Center for Mineral Metabolism and Clinical Research, University of Texas Southwestern Medical Center, Dallas, TX, 75390, USA; Department of Internal Medicine, University of Texas Southwestern Medical Center, 5323 Harry Hines Blvd, Dallas, TX, 75390, USA

## Abstract

The uterus is a hollow, fibromuscular organ involved in physiologic processes such as menstruation and pregnancy. The content and organization of extracellular matrix constituents such as fibrillar collagen dictates passive (non-contractile) biomechanical tissue function; however, how extracellular matrix composition and biomechanical function change with age in the uterus remains unknown. This study utilizes Raman spectroscopy coupled with biaxial inflation testing to investigate changes in the murine uterus with age (2-3 months, 4-6 months, 10-12 months, and 20-24 months). Linear and toe moduli significantly decreased with reproductive aging (2 to 12 months); however, moduli increased in the oldest age group (20-24 months). The optical signature of combined elastin and collagen content was significantly higher in the oldest group (20-24 month), while the glycogen contribution was the highest in the 2-3 month murine uterus. The presented workflow couples biaxial inflation testing and Raman spectroscopy, representing a critical first step to correlating biomechanics and optical signatures in the aging uterus with the potential for clinical translation. Further, this study may provide critical compositional and structure-function information regarding age-related uterine disorders.

## Introduction

The uterus is a dynamic organ that grows and remodels in response to varying biochemical and biomechanical cues throughout a woman’s lifespan such as menstruation and pregnancy [1-6]. The composition and arrangement of cells and the non-contractile or passive components of the extracellular matrix (ECM) dictate uterine function. The primary load-bearing components of the uterus are collagen fibers, elastic fibers, and smooth muscle cells [7, 8]. Uterine smooth muscle cells work with the non-contractile (passive) components of the matrix to maintain function and stability throughout physiologic processes [4, 9]. Contributing to passive biomechanical function, collagen fibers impart tensile strength to the tissue while elastic fibers permit stretching and recovery [8, 10, 11]. Cells within soft tissues, such as fibroblasts, remodel their ECM; however, disruptions in this process with aging contribute to deleterious remodeling and diminished biomechanical function [12-15]. Previous work quantified collagen fibrosis in the murine uterus wherein collagen deposition increased with advanced age [16]. Aging also reduced contractile function in the murine and rat uterus [17, 18]. However, how the passive biomechanical and compositional properties of the murine uterus change with age are not fully understood. Elucidating these changes may provide insights into age-related reproductive dysfunction and its adverse physiological impacts, such as increased risk of prolonged labor with advanced maternal age [17].

Prior work in uterine biomechanics quantified uniaxial properties in human tissue [19-23] and animal tissue [24-26]. While these results serve as a useful baseline for establishing biomechanical properties of the uterus, the uterus is loaded multiaxially in the body due to neighboring organs, supporting ligaments, intraabdominal pressure, and gravity [27, 28]. Biaxial inflation methods retain cylindrical geometry and native cell-ECM interactions and are used in the vasculature [29, 30], vagina [12, 13, 31-35], cervix [36], and utero-cervical complex [37]. Biomechanical testing provides insight into the function of the tissue, but the microstructural and compositional changes driving the functional behavior need quantification.

Towards this end, Raman spectroscopy (RS) is a nondestructive and noninvasive optical technique that utilizes incident light to extract the molecular fingerprint of a sample. These molecular fingerprints arise from vibrational bonds of specific compositional components in a sample. Prior work in colon [38-40], cervix [41-43], breast [44, 45], and other tissues [46] demonstrates the sensitivity of RS to compositional changes in biological applications. RS is an effective tool with the capability to identify optical biomarkers for otherwise difficult to distinguish pathologies. Inflammatory bowel disease (ulcerative colitis and Crohn’s disease) has overlapping symptoms and pathologies, commonly resulting in misdiagnosis; however, RS successfully detected disease-specific biomarkers in the colon, markedly improving differentiation [38-40]. Additionally, RS optically quantified significant changes in collagen and elastin contributions during gestation in the cervix, proving useful in predicting preterm birth [41-43]. Extending this approach to combine RS with biomechanical function in the uterus will provide a comprehensive understanding of how structure and function of the uterus changes with age.

Mouse models are commonplace in reproductive research due to their quick breeding cycles, relatively economical upkeep, and short lifespan [5, 6, 37, 47]. While not a complete replacement for human tissue, mice are a useful starting point for identifying structure-function relationships and as cost-effective model systems to quantify changes with age [12, 13]. Although the mouse and human uteri differ in shape, they both consist of an inner epithelial-lined endometrium and muscular myometrium layers [48]. Further, the well documented age correlations between these species in addition to the difficulty studying extended, progressive age-related disorders in humans make mice an ideal model of uterine aging [49].

Therefore, the objective of this study was to quantify the passive biomechanical properties and composition of the nulliparous murine uterus with age using biaxial inflation protocols and RS. We hypothesize that distensibility will increase with reproductive age, corresponding to a decrease in elastin and collagen content [17, 50]. If successful, these results will provide fundamental biomechanical and compositional information on the aging uterus, which may have clinical implications for pregnancy complications associated with advanced maternal age. Additionally, this study may provide key insights into uterine changes with reproductive senescence. Finally, this study will serve as a confirmation of the power of coupling RS and biomechanics with implications for clinical translation for other uterine conditions such as fibroids and endometriosis.

## Materials and Methods

### Animal Use and Specimen Preparation

This study used 32 female nulliparous wildtype C57BL/6 mice (UTSW IACUC approved). All animals were housed in a 12-hour light/dark cycle with access to standard chow. Mice were divided into four groups according to age: 2-3 months, 4-6 months, 10-12, and 20-24 months of age (n=8/group) (Fig. 1A). Mice were euthanized via cervical dislocation, and uterine horns were dissected from the reproductive tract (Fig. 1B) and frozen at −20°C. Prior to testing, the tissue was defrosted for one hour at 4°C. One freeze-thaw cycle has no effect on passive mechanical properties [51]. One uterine horn was randomly allocated to biomechanical testing, while the other was used for Raman Spectroscopy.

**Figure 1.**
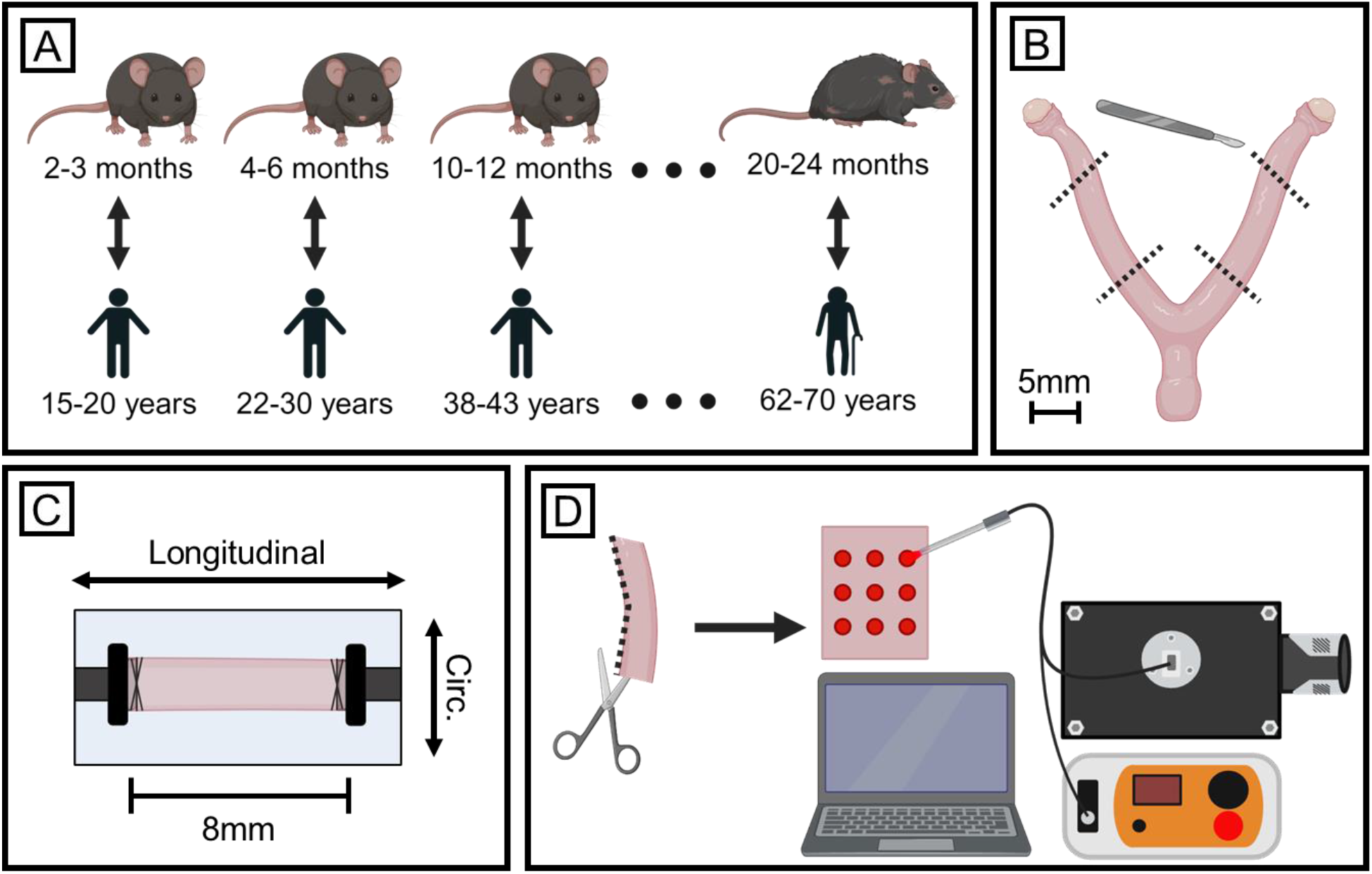
Panel of correlative biomechanics and optical spectroscopy workflow developed for assessment of murine uterine tissues. **(A)** Age groups of the mice in months with their corresponding age in human years [49]. **(B)** Murine uterus dissection. Cuts were made at the dashed lines, and the uterine horns were randomly allocated for biaxial inflation testing or Raman spectroscopy. **(C)** Cannulated uterine horn for biaxial inflation testing. **(D)** A longitudinal cut was made to expose the lumen of the uterine horn, and nine regions of interest were probed.

### Biomechanical Testing

A transverse cut was made 8 mm above the uterine bifurcation point for biomechanical testing (Fig. 1B). The tissue was cannulated onto a biaxial inflation device (Danish MyoTechnologies, Aarhus, Denmark) with 0.75 mm cannulas (Fig. 1C). Three 6-0 silk sutures on each end secured the tissue in place and the organ bath was filled with calcium-free Kreb’s Ringer Buffer (KRB) at 37°C. Due to tissue retraction after excision, the tissue was extended longitudinally to the point wherein tension nor compression was experienced, restoring the unloaded length [31, 52]. Following a 30-minute equilibration period, the uterus was circumferentially preconditioned for five cycles of cyclic increasing and decreasing pressure from 0 to 140 mmHg. Following the theory that tissues seek to preserve energy *in vivo*, the tissue was extended to the length where force remained constant over a range of increasing pressure, restoring the estimated physiologic length [13, 53, 54]. Five more cycles of circumferential preconditioning were performed from 0 to a maximum pressure of 200 mmHg to minimize hysteresis and achieve consistent and repeatable measurements [37]. For longitudinal preconditioning, the tissue was extended longitudinally from −1% below the estimated physiologic length to 1% above at 1/3 of the maximum pressure (67mmHg). The tissue equilibrated for 10 minutes followed by re-establishing the unloaded geometry. Three cycles of pressure-inflation testing (P=0-200 mmHg) were performed at the physiologic length, −1% and +1% of the physiologic length. Force length testing was performed by cycling between −1% and +1% of the physiologic length at pressures of 10, 67, 133, and 200 mmHg [37]. The last loading cycle of each test was used for data analysis. Following biomechanical testing, a 0.5 mm thick ring was cut transversely where outer diameter tracking was performed [55]. The ring was imaged in KRB using a stereo microscope (Stereozoom S9i, Leica Instruments, Singapore). The length of 25 transmural lines were traced circumferentially in ImageJ (National Institutes of Health, Bethesda, MD, USA) and averaged to quantify the estimated unloaded thickness. This process was repeated two more times, and the final three technical replicates were averaged to determine the thickness of the tissue [37, 55].

### Raman Spectroscopy

This study utilized a back-illuminated, deep-depletion charge-coupled device, thermoelectrically cooled to −70°C (PIXIS: 400BRX, Teledyne Princeton Instruments, Princeton, NJ, USA) in conjunction with a high throughput spectrometer (HT3-SPEC-785, EmVision LLC, Loxahatchee, FL, USA). A custom 2.1mm outer diameter Raman probe with seven collection fibers arranged around one excitation fiber (EmVision LLC, Loxahatchee, FL, USA) was connected to the spectrometer and a 785nm laser (I0785MM0350MS-USB, Innovative Photonic Solutions Inc., Plainsboro, NJ, USA) for data collection. Before each experiment, the laser was calibrated with a power meter (PM16-121, ThorLabs, Newton, NJ, USA) to deliver 80mW from the distal tip of the fiber optic probe. Spectra of a neon-argon lamp were acquired for absolute wavenumber (wavelength) calibration, while measurements of acetaminophen and naphthalene samples were taken to calculate the exact Raman shift (relative wavenumber axis). Measurements of a calibrated National Institute of Standards and Technology (NIST) Tungsten and Krypton lamp (SL1-CAL, StellarNet Inc., Tampa, FL, USA) and glass Standard Reference Material (SRM2241, NIST, Gaithersburg, MD, USA) enabled spectral response correction of the acquired Raman spectra. The uterine horn was opened longitudinally to expose the endometrium where nine regions of interest were identified in a grid formation (Fig. 1D). Ten experimental replicates were acquired for each point on the grid at an integration time of 250ms. A Savitzky-Golay filter smoothed the spectral noise, and the biological fluorescence was subtracted using an established, modified polynomial fit algorithm [56]. Finally, the spectra were normalized to the intensity of the phenylalanine peak (Supp. Fig. 1).

### Biomechanical Data Analysis

For biomechanical analysis, the uterus was assumed to be a uniform, incompressible cylinder. Equation 1 calculated the unloaded volume (V) using unloaded outer radius (*R*_*o*_), estimated unloaded thickness from ring measurements (*H*), and unloaded length (*L*).

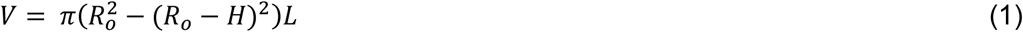

Enforcing the assumption that the uterus behaves as an incompressible soft tissue, Equation 2 determined the deformed inner radius (*r*_*i*_) throughout the experiment based on the deformed length (*l*) and outer radius (*r*_*o*_).

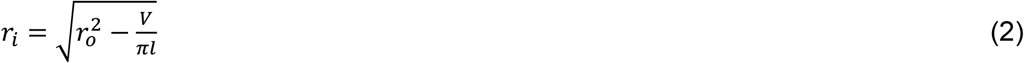

Equations 3 and 4 quantified the circumferential (*σ*_*θ*_) and longitudinal (*σ*_*z*_) Cauchy stresses [13, 54].

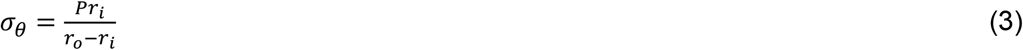

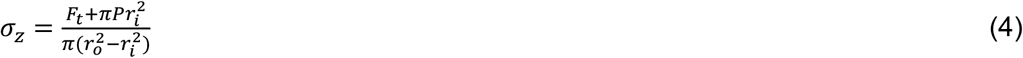

*P* is intraluminal pressure and *F*_*t*_ the force measured by the force transducer. Circumferential stretch (*λ*_*θ*_) and longitudinal stretch (*λ*_*z*_) were calculated by Eq 5 and 6, respectively, using the deformed length (*l*).

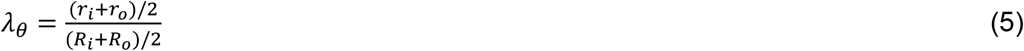

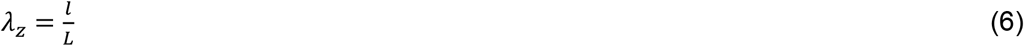

To determine the toe and linear regions of the stress-stretch curves, a bilinear curve fit MATLAB *lsqcurvefit* function was applied (Supp. Fig. 2) The slope of each region quantified their respective modulus, a measure of resistance to deformation [37, 52].

### Raman Spectroscopy Data Analysis

A MATLAB code fit Gaussian curves under the Amide III (1265 cm^-1^), Amide I (1670 cm^-1^), and protein/lipid (1440-1450 cm^-1^) features to extract the intensity and width of these peaks and relate them to protein secondary structure (Fig. 2) [43, 57, 58]. Two peak ratios (850 cm^-1^/830 cm^-1^ and 1304 cm^-1^/1265 cm^-1^) were calculated using the intensity values extracted from the Gaussian fits for additional compositional information. Spectra of pure components collagen (bovine Achilles), glycogen (bovine liver), elastin (bovine neck ligament), cholesterol, hyaluronic acid (Sigma-Aldrich, Co., St. Louis, MO), and adipose tissue (murine C57BL6 abdomen) were acquired as inputs for mathematical modeling of uterine composition. The murine abdominal adipose tissue most accurately represents the multifaceted biological fat; however, it was compared to four pure lipids (glyceryl trimyristate (palm plant), glyceryl trioleate (sunflower plant), l-α-phosphatidylcholine (egg yolk), and palmitic acid (palm oil)) and confirmed to correlate significantly with l-α-phosphatidylcholine (Supp. Fig. 3, Supp. Fig. 4). Previous studies of the aorta and cervix used these components [41-43, 59]. A Spearman correlation analysis of the six pure components accounted for the assumption of input variable independence in the non-negative least squares (NNLS) algorithm (ρ > 0.70 indicated highly correlated spectra) (Supp. Fig. 3). Elastin and collagen signals were combined due to their collinearity and the relatively miniscule elastin content in the uterus [60, 61]. A Lawon-Hanson implementation of the NNLS algorithm was performed with the “nnls” package in R Studio to determine the contribution of these pure component signals to the overall uterine signals (Fig. 3) [62].

**Figure 2.**
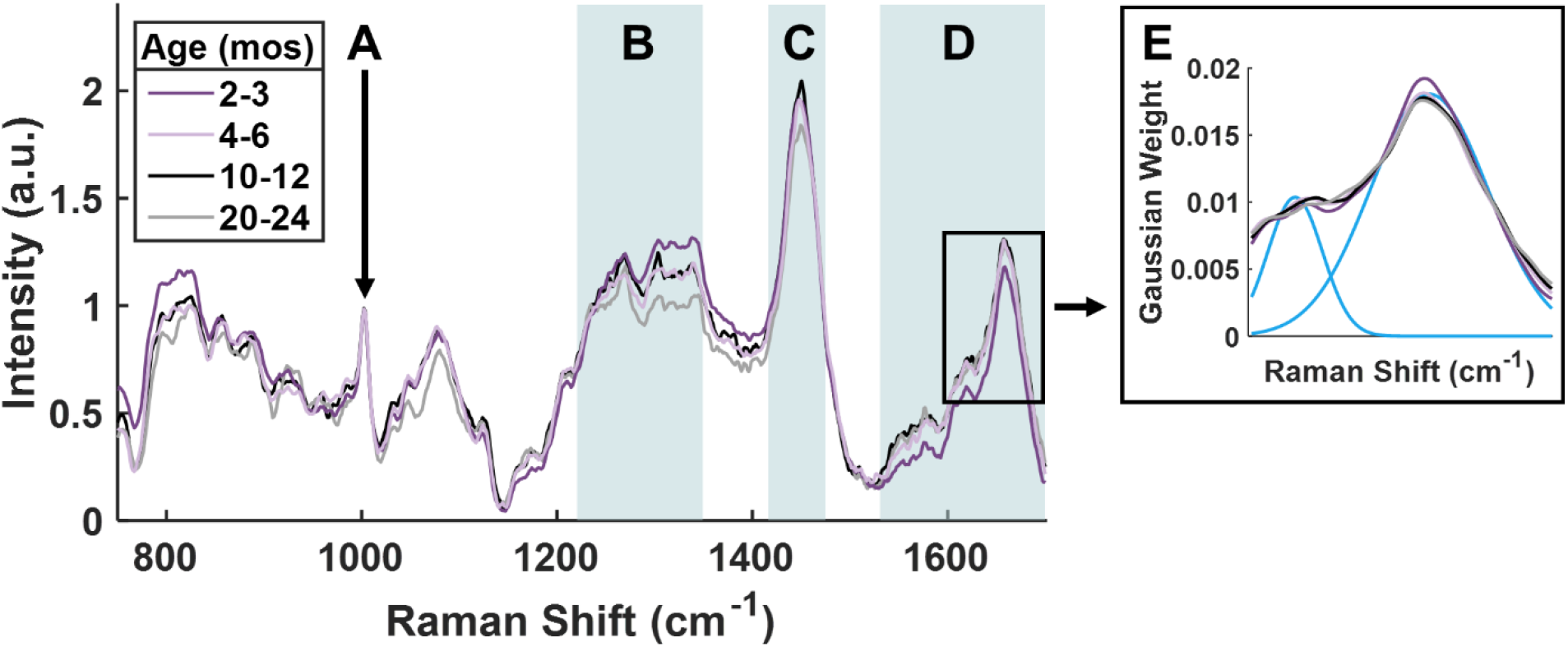
Averaged Raman spectra per age group with indicated regions of interest for peak ratios and feature extraction: **A**. Phenylalanine peak. **B**. Amide III. **C**. Lipid-protein peak. **D**. Amide I. **E**. Example Gaussian fit of the Amide I 1609 cm ^-1^ shoulder and 1657 cm^-1^ peak.

**Figure 3.**
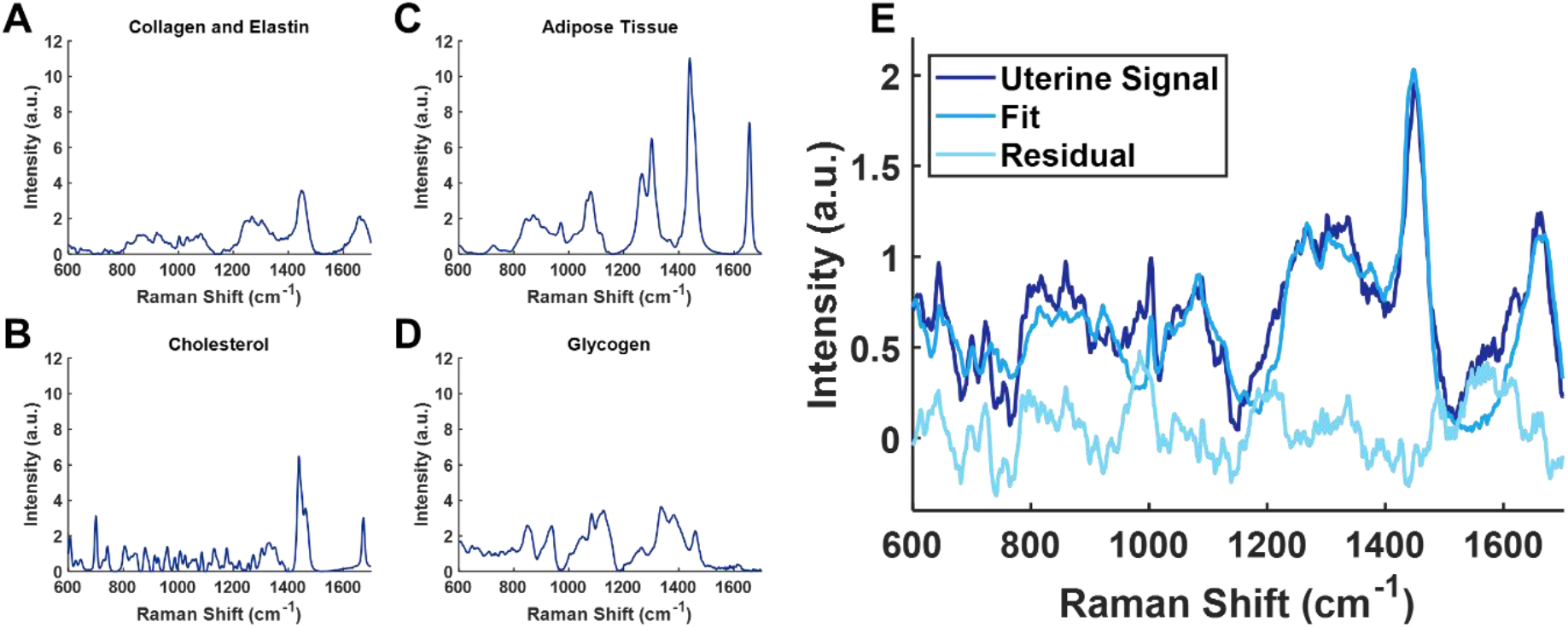
Individual spectra for model components **(A)** collagen and elastin, **(B)** cholesterol, **(C)** mouse fat, and **(D)** glycogen. **(E)** NNLS analysis of the uterine spectra from a 4-6 month old mouse plotted against the calculated fit (y) and the residuals (ε) [Eq. 7].

The NNLS algorithm is a supervised, regression-based model that uses pure spectral inputs (*X*), their calculated coefficients (β), and an error term (ε) to fit a curve (y) to the original measured signal [Eq. 7].

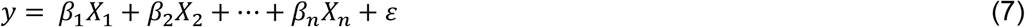

### Statistical Analysis

All statistical analyses were performed in RStudio (4.3.1). To assess the homogeneity of variances and normality of each measure, Levene and Shapiro-Wilks tests were performed, respectively. A one-way ANOVA test evaluated differences between geometry (estimated physiologic length, outer diameter, thickness) with respect to age and Tukey’s post hoc test followed. A one-way ANOVA with Tukey’s post hoc test quantified the differences in the toe and linear moduli with respect to age. For non-normally distributed data, a nonparametric Kruskal-Wallis or Scheirer Ray Hare (SRH) test paired with Dunn’s post hoc test was used. For compositional comparisons (NNLS coefficients, peak widths, and ratios), a multinomial regression was applied using the “nnet” package to determine the coefficients’ impact on the murine uterine age with respect to the nine technical replicates for each biological replicate [63]. From the multinomial model, the coefficients were extracted for each group and transformed into z-values, where the cumulative density function was utilized to calculate p-values.

### Results

#### Geometry

Thickness significantly increased (Fig. 4A) in the 20-24 month uterus compared to the 2-3 month group (p=0.04). While not significant (p=0.23), estimated physiologic length decreased with age (Fig. 4B). Outer diameter at 1/3 maximum pressure increased from 2-3 months to 4-6 months, then decreased in 10-12 months (Fig. 4C). The outer diameter increased again from 10-12 months to 20-24 months.

**Figure 4.**
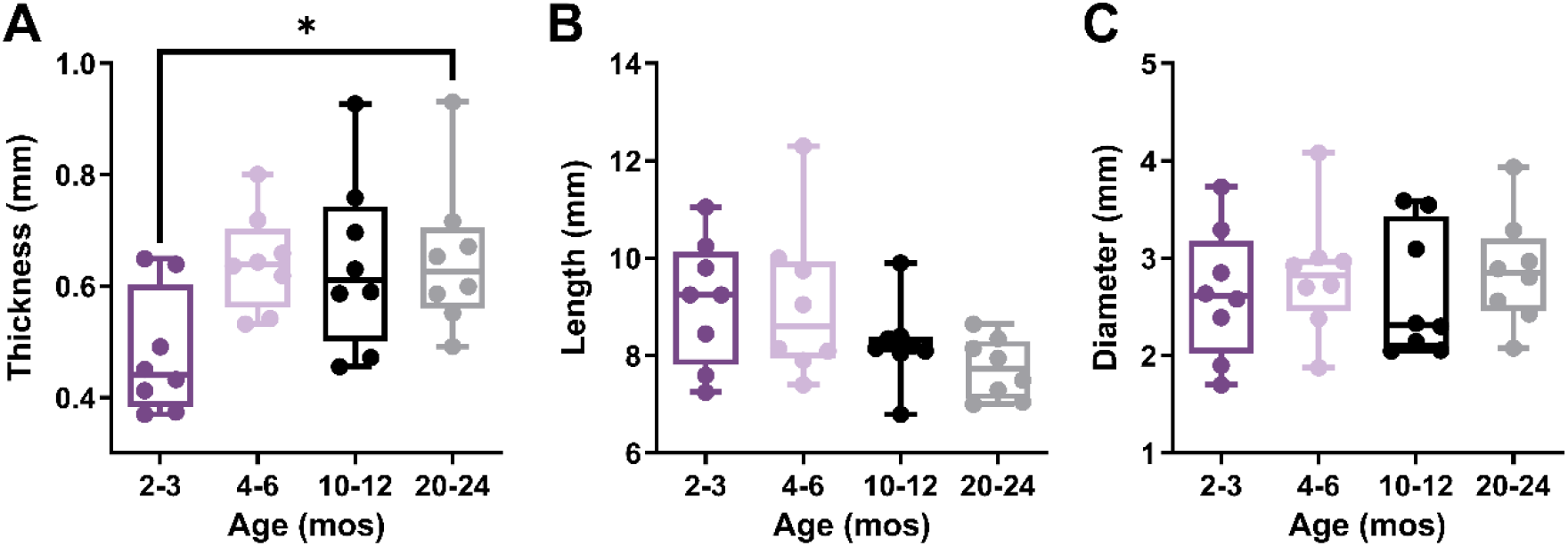
Thickness **(A)**, estimated physiologic length **(B)**, and outer diameter **(C)** for the 2-3 month (dark purple), 4-6 month (light purple), 10-12 month (black), and 20-24 month (grey) uterus. No significant differences were identified in estimated physiologic length (p=0.23) or outer diameter (p=0.75). Thickness significantly increased in the 20-24 month uterus compared to the 2-3 month (p=0.04). ^*^p<0.05 denotes statistical significance, n=8/group.

#### Biomechanics

The rightward shift of the 4-6 month and 10-12 month uterine stress-stretch curves from the 2-3 month group showed increased distensibility (Fig. 5). The stress-stretch curve of the 20-24 month group shifted leftward, indicating decreased distensibility compared to the 4-6 month and 10-12 month groups. The 20-24 month group was more distensible than the 2-3 month group. The tissue in both the circumferential (Fig. 5A) and longitudinal (Fig. 5B) directions showed similar trends.

**Figure 5.**
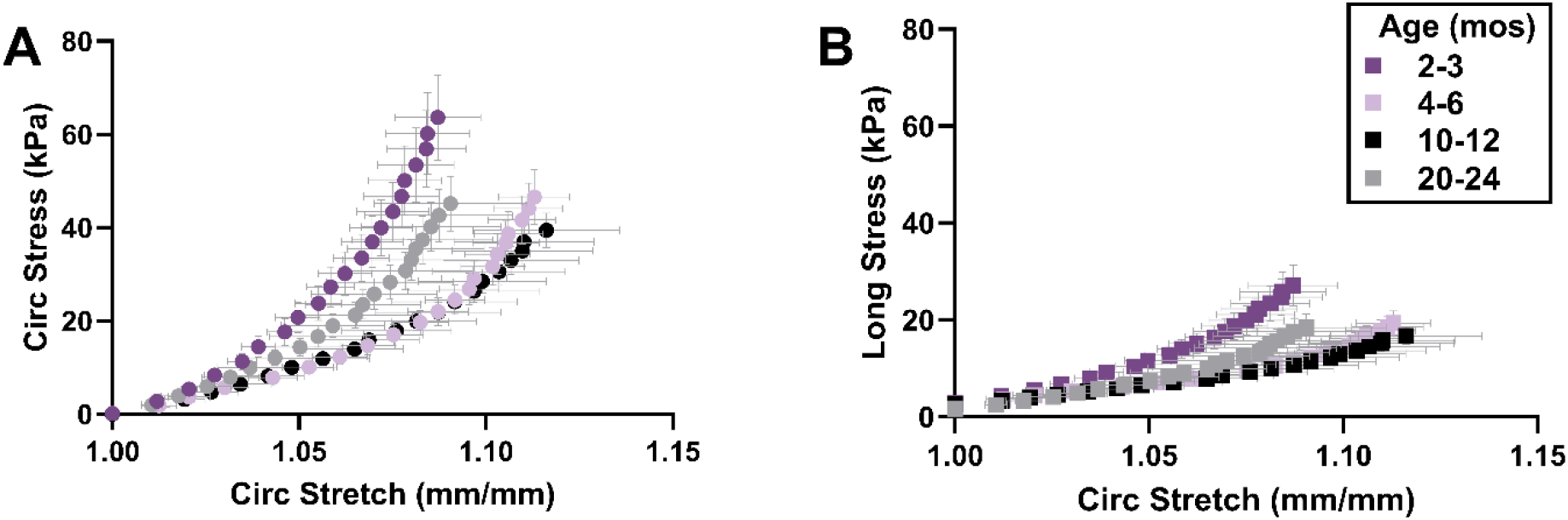
Circumferential **(A, circles)** and longitudinal **(B, squares)** stress-stretch curves for the 2-3 month (dark purple), 4-6 month (light purple), 10-12 month (black), and 20-24 month (grey) uterus. Distensibility increased with reproductive aging (4-6 and 10-12 months) compared to the 2-3 month group in both directions. Data is reported as mean ± standard error of mean, n=8/group.

Significant differences in the circumferential toe modulus were identified between age groups (p=0.015) (Fig. 6A). Circumferential toe modulus significantly decreased in the 4-6 (p=0.05) and 10-12 month (p=0.03) uterus compared to the 2-3 month uterus. Age significantly affected circumferential (p=0.01) and longitudinal (p=0.04) linear moduli (Fig. 6B). Circumferential linear modulus significantly decreased in the 10-12 month uterus compared to the 2-3 month (p=0.009). Longitudinal linear modulus also significantly decreased in the 10-12 month uterus compared to the 2-3 month (p=0.03). No other significant comparisons were detected.

**Figure 6.**
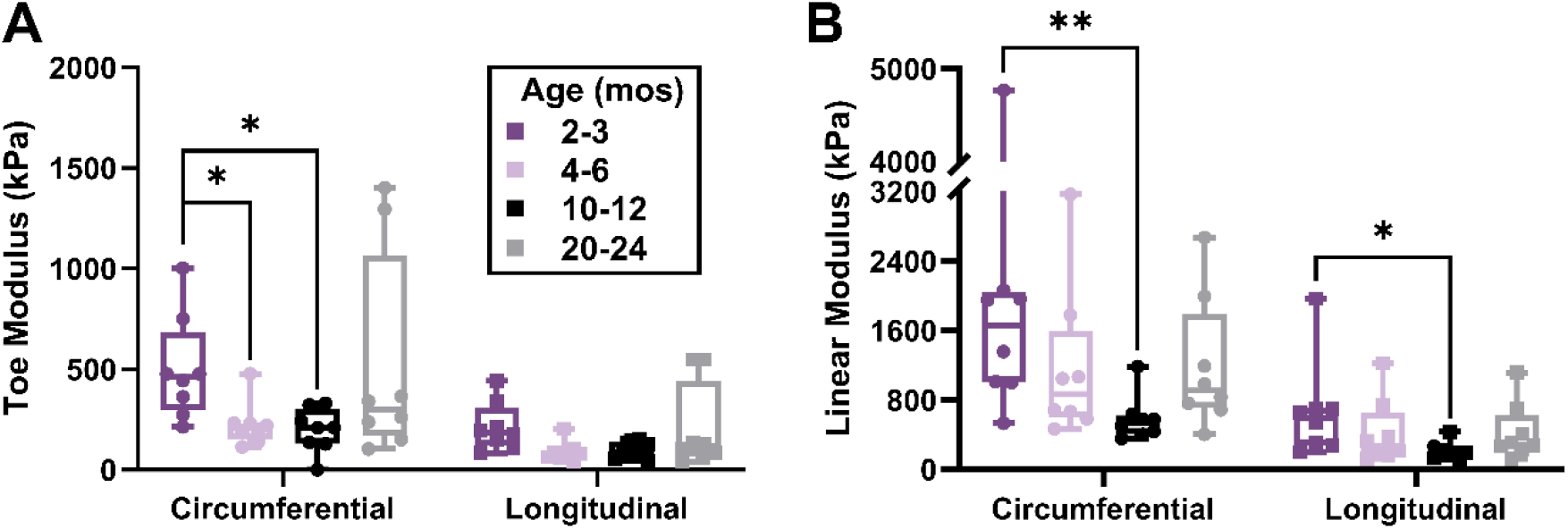
Toe **(A)** and linear **(B)** modulus in the circumferential (circles) and longitudinal direction (squares) for the 2-3 month (dark purple), 4-6 month (light purple), 10-12 month (black) and 20-24 month (grey) uterus. For the toe modulus, circumferential modulus significantly decreased in the 4-6 (p=0.05) and 10-12 month (p=0.03) uterus compared to the 2-3 month group. The 10-12 month uterus had a significantly lower linear modulus than the 2-3 month group in the circumferential (p=0.009) and longitudinal (p=0.03) directions. No other significant differences were identified. ^**^p<0.01, ^*^p<0.05 denotes statistical significance, n=8/group.

#### Raman Spectroscopy

A multinomial regression model identified significant differences between the glycogen, elastin and collagen, and cholesterol NNLS scores (Fig. 7). The 2-3 month glycogen score was significantly higher than the other age groups (p<0.0001) (Fig. 7A). Conversely, the elastin and collagen NNLS score was elevated in the 20-24 month uterus compared to the 2-3 month (p=0.003), 4-6 month (p=0.02), and 10-12 month (p=0.04) age groups (Fig. 7B). Finally, the cholesterol contribution significantly decreased in the oldest age group with respect to the 2-3 month (p=0.02), 4-6 month (p=0.002), and 10-12 month (p<0.0001) uterus (Fig. 7C). The 10-12 month cholesterol NNLS scores also differed significantly from the 2-3 month group (p=0.01).

**Figure 7.**
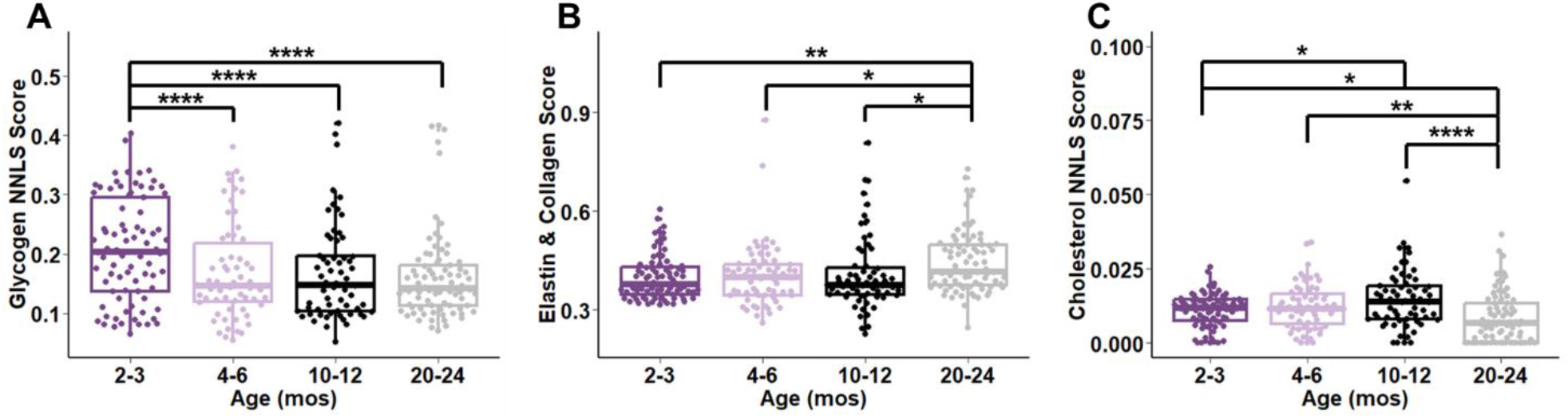
Glycogen **(A)**, elastin and collagen **(B)**, and cholesterol **(C)** NNLS scores for the 2-3 month (dark purple), 4-6 month (light purple), 10-12 month (black), and 20-24 month (grey) uterus. Significant differences were identified for each of the three pure components with respect to uterine age. ^****^p<0.0001, ^***^p<0.001, ^**^p<0.01, ^*^p<0.05 denotes statistical significance, n=8/group.

A multinomial regression model identified significant differences between the width of the Amide I peak and the β-sheet to random coil ratio. The youngest age group had a significantly smaller Amide I width compared to the other age groups (4-6 month, p=0.0002, 10-12 month, p=0.004, 20-24 month, p<0.0001) (Fig. 8A). Amide I width was also significantly lower in the 10-12 month uterus with respect to the 20-24 month age group. The Amide III β-sheet to random coil ratio was significantly lower in the 4-6 month group when compared to 10-12 month (p=0.01) and 20-24 month (p=0.002) (Fig. 8B).

**Figure 8.**
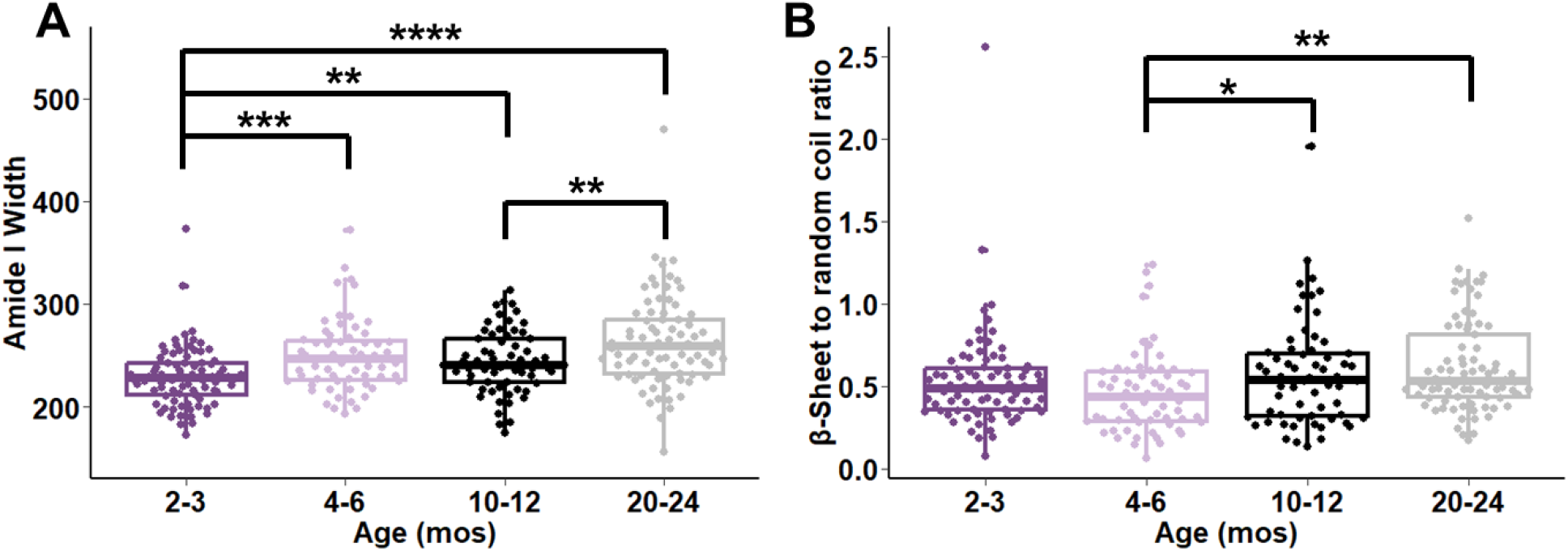
Amide I width **(A)** and Amide III β-sheet to random coil ratio **(B)** for the 2-3 month (dark purple), 4-6 month (light purple), 10-12 month (black) and 20-24 month (grey) uterus. The youngest age group showed the smallest Amide I width when compared to other uterine spectra (4-6 month, p=0.0002, 10-12 month, p=0.004, 20-24 month, p<0.0001). The β-sheet to random coil ratio increased with age from the 4-6 month to the 10-12 month (p=0.01) and 20-24 month (p=0.002) groups. ^****^p<0.0001, ^***^p<0.001, ^**^p<0.01, ^*^p<0.05 denotes statistical significance, n=8/group.

## Discussion

This study presents a comprehensive workflow for correlative analysis of passive (non-contractile) biomechanical function and composition of the isolated murine uterus using biaxial inflation and Raman spectroscopy methods. This report, to the best of our knowledge, represents the first research integrating both optical spectroscopy and biomechanics for complementary composition – functional analysis in uterine tissues. Additionally, this study applied the combined method to determine biomechanical and compositional changes of the murine uterus as a function of age, demonstrating complementary shifts in properties as mice progress towards reproductive maturity that differ from mice aging towards senescence.

### The uterus was more distensible with reproductive aging

Uterine distensibility increased (Fig. 5) and modulus decreased (Fig. 6) with reproductive aging (2 to 12 months). The increased cholesterol contribution (Fig. 7C) with reproductive aging may contribute, in part, to the evolving biomechanical function. Cholesterol is a hormone-dependent lipid that increases stiffness in human tendons, thus reducing functionality [64]. Additionally, it is a critical precursor to local steroid hormone production in the reproductive system. Estrogen and progesterone regulate collagen and elastic fibers in the cervix, subsequently dictating mechanical function [50, 65]. Cholesterol rapidly increases in women between the ages of 40-64 years and inhibits uterine contractility [66, 67]. Because the uterus is a hormonally driven organ, decreasing progesterone and estrogen levels with age may impact the changes in elastic and collagen fiber content [68, 69]. Altogether, age-related changes in cholesterol content may impact uterine passive mechanics, leading to complications in pregnancies of advanced maternal age

We hypothesized that total elastin and collagen content would decrease with age; however, Raman results did not confirm this (Fig. 7B). Separate changes in collagen and elastic fiber quantities or microstructural components may contribute to increased distensibility (Fig. 5) and decreased toe and linear modulus (Fig. 6) with age. Within vasculature, loss of elastic fiber integrity and collagen remodeling increases vascular stiffness with age [70]. Reduced elastic fiber components such as elastin, fibrillin, and fibulins and decreased collagen fiber undulation may give rise to age-related biomechanical changes [71]. In the murine vagina, elastase treatment significantly decreased contractility and increased modulus in 2-3 month and 4-6 month groups [12, 13]. In the 7-9 month and 10-14 month vagina, elastase did not significantly affect the mechanical response suggesting that younger mice may have a higher amount of intact, functional elastic fibers than the older groups [12, 13]. Further, hydroxyproline, an amino acid found in collagen, increases with age in the cervix [72, 73]. What fibrous protein quantities and microstructural components change with age in the uterus remains unknown. Future studies are needed to determine uterine fiber quantities and related microstructural components with age to provide insights into age-related uterine remodeling and their relations to adverse maternal outcomes in pregnancy.

While the microstructural mechanisms dictating changes in passive uterine behavior are not fully understood, changes in passive mechanics may coincide with or contribute to decreased contractile behavior. This may, in part, contribute to increased risk of labor dystocia with age [74]. Mice at eight months old have longer gestation times, extended periods of labor, and fewer pups than mice at three months old [17]. Further, uterine contractile function decreases with age and requires external agonists to reach maximum potential [18]. During pregnancy and parturition, the passive matrix components and smooth muscle cells remodel, resulting in altered mechanical function to accommodate fetal growth and permit safe delivery [1, 2, 5, 23, 42]. Age-related ECM and smooth muscle cell remodeling in the nulliparous state may lead to complications during pregnancy with advanced maternal age. Further studies are needed to better understand the relationship between uterine contractile function and ECM composition with age.

### Reproductive senescence led to tissue stiffening and potential fibrosis

While uterine toe and linear modulus decreased with reproductive aging, both moduli increased in the 20-24 month group which is undergoing reproductive senescence (Fig. 6). The increased elastin and collagen content (Fig. 7B) and subsequently increased tissue thickness observed herein (Fig. 4A) may contribute. Furthermore, Amide I significantly increased with senescence (Fig. 8A), which may represent decreased collagen organization. In agreement with these findings, collagen deposition in the murine uterus significantly increases in the 18-month versus 3-month group [16]. Collagen deposition and fibrosis also occur in the murine and human ovary with age and correlate with increased ovarian stiffness [75]. We hypothesize that the simultaneous accumulation and increased disorganization of fibers may explain the increased modulus in the senescent group and is a natural part of aging in the uterus.

In agreement with prior work, this study identified a significant decrease in cholesterol with senescence (Fig. 7C). Cholesterol levels in women decline with age later in life. Cholesterol depletion in a mink model of cardiovascular disease is linked to autophagy and arterial stiffening [76]. The impact of cholesterol on the aging uterus is unknown; however, reproductive senescence correlates to uterine atrophy [77]. In contrast to the decreased cholesterol contribution in the 20-24 month murine uterus, the linear and toe moduli increased. These findings support the previously mentioned hypothesis that a relationship between optical cholesterol content and uterine passive mechanics may exist.

Further, endometrial cancer is prominent in senescent age groups [78]. While the cause of endometrial cancer is not understood, fibrosis and changes in mechanical properties may influence abnormal cellular growth [79]. Future studies should consider age-related remodeling of the uterus when investigating endometrial cancer.

In addition to content and microstructural features such as organization, protein conformation also contributes to tissue biomechanical properties [80-83]. However, the effect of protein secondary structure on uterine biomechanical function remains unclear. The β-sheet to random coil ratio was not significant between the youngest and oldest groups, but the 20-24 month murine uterus had a significantly higher ratio than the 4-6 month and 10-12 month groups (Fig. 8C). Altered distributions of protein secondary structures in the 20-24 month group (Fig. 5B) may explain why distensibility and modulus 20-24 month approached levels of the 2-3 month group [82]. This may imply that protein conformation, in part, dictates uterine passive function.

### Murine uterine glycogen changed with age

Glycogen content was significantly higher in the 2-3 month murine uterus (Fig. 7A). These findings are consistent with prior literature in the vagina of an ewe model of menopause [84]. Decreasing glycogen content is associated with vaginal wall thinning and atrophy which is consistent in post-menopausal women [84]. However, it remains unknown how glycogen concentration impacts uterine biomechanics. Glycogen is a critical component of the uterine endometrium and is directly associated with fertility; however, the uterus cannot store glycogen [85]. Alternatively, dysregulated glycogen metabolism in the uterus contributes to autophagosome buildup in mice [86], wherein autophagy in rat epithelial uterine tissue results in atrophy and decreased distensibility [87]. An association with decreased glycogen content and uterine moduli may exist, contributing to increased incidence of spontaneous abortion, infertility, and prolapse with advanced maternal age and reproductive senescence. However, further research is needed.

### Changes in biomechanical function and composition differed between reproductive organs

Distensibility followed a similar trend shown in the C57 murine cervix from 3 to 8 months of age [17]. However, results in the uterus were opposite from results quantified in the CD-1 murine vagina: from 2 to 14 months, distensibility progressively decreased while modulus increased [12]. Differing results between reproductive organs emphasizes the need to assess biomechanical properties when the organs are conjoined for physiologic and clinically translatable results.

This study was not without limitations. The reproductive cycle, or the estrous cycle in rodents, was not controlled in this study which may contribute to variability in results [88]. Glycogen and progesterone are highly related, indicating that the lack of consistent estrus cycling may increase the glycogen NNLS score variability [88]. However, there are no reported significant differences in passive mechanics in the utero-cervical complex or rat vagina throughout the estrous cycle [37, 89]. Further, while the short life cycles of mice are advantageous for age related studies, hormonal changes differ in mice compared to humans with age. Humans have a cyclic menstrual cycle that lasts between 10-40 years [90-92]. Towards the end of a woman’s fertile years, the transition to menopause begins wherein ovarian function is lost, circulating estrogen declines, and the cyclic menstrual cycle becomes irregular and eventually ceases [93, 94]. Unlike humans, mice continue to cycle throughout middle age and do not experience a menopausal phase. Cycle frequency begins to decline around 10-12 months of age and mice become acyclic between 18-24 months of age when they are considered “fully aged” [47, 95, 96]. In animal models of human reproductive aging, it is common to perform an ovariectomy to induce menopause [97]. Thus, the 20-24 month murine group is thought to best represent reproductive senescence in our study with the acknowledgment that these mice were not models of menopause. While not an exact representation of human aging, this study is a crucial first step at understanding the impact age has on biomechanical and compositional changes in the uterus [47, 91, 92, 94-96].

In summary, this study quantified biomechanical and compositional changes with uterine aging through biaxial inflation and optical spectroscopy analysis methods for the first time. Identifying biomechanical and compositional changes provides crucial information on how the uterus remodels with reproductive senescence as well as potentially offering insight into pregnancy complications with advanced maternal age. Finally, this work represents a critical first step to establishing a workflow correlating biomechanics and optical spectroscopy with implications of clinical translation into identification of uterine pathophysiology.

## Acknowledgements

The authors would like to acknowledge Haolin Shi for animal husbandry. Additionally, this research would not be possible without the feedback and moral support of Lily Buchanan and Jessica Baas. This work is funded, in part, by NSF CMMI 2053851 (De Vita, Abramowitch, Miller, Myers), UT Rising STARs Award (Pence), AHA Second Century Early Faculty Independence Award 23SCEFIA1156385 (Pence) and AAOGF/Wellcome Boroughs Career Development Award (Florian-Rodriguez).

## Supplementary Figures

**Supplemental Figure 1.**
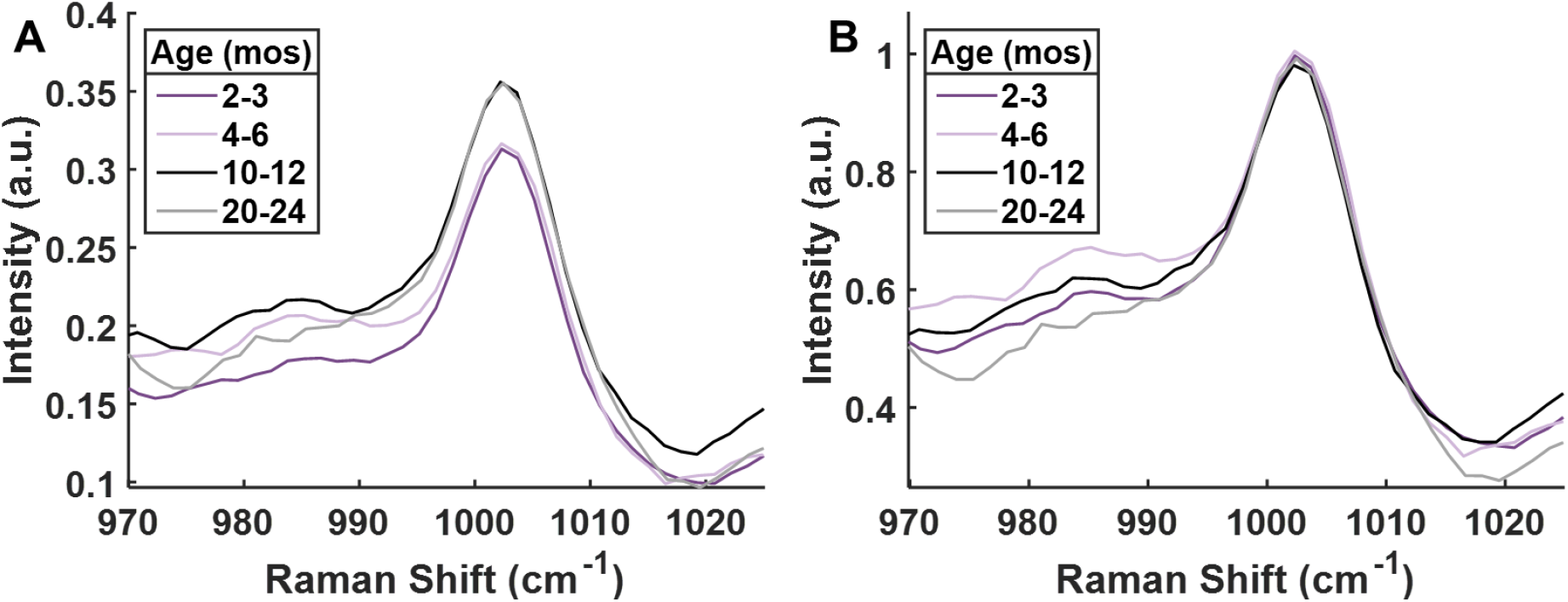
Averaged spectra from each age group before normalization to the phenylalanine peak at 1003 cm^-1^ **(A)** and after **(B)**.

**Supplemental Figure 2.**
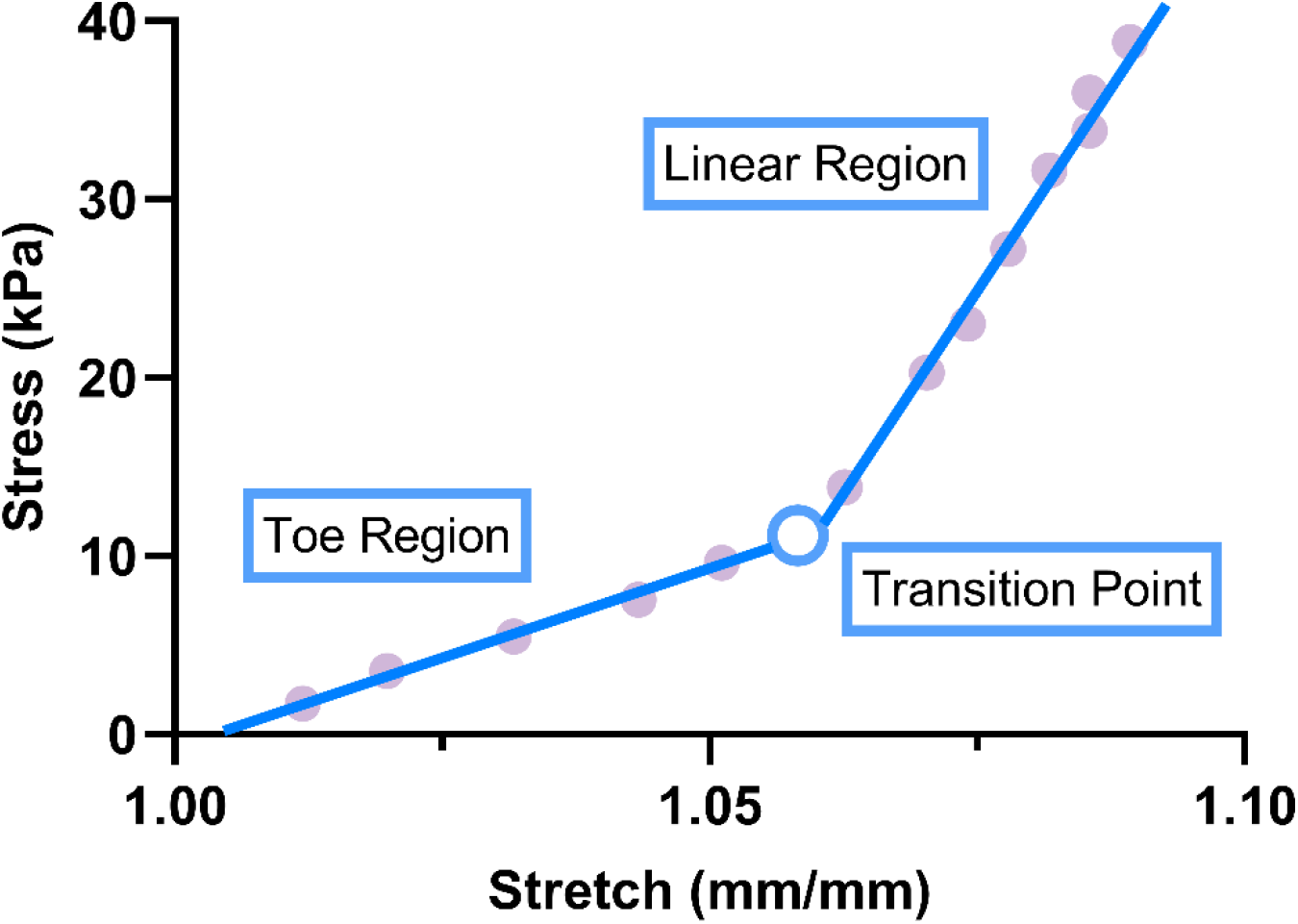
Schematic of a bilinear curve fit (blue line) to the toe and linear region of the circumferential stress-stretch curve for a 4-6 month old (light purple circles) murine uterus. The MATLAB function *lsqcurvefit* quantified toe and linear moduli as the slope of their respective region and the transition point [52].

**Supplemental Figure 3.**
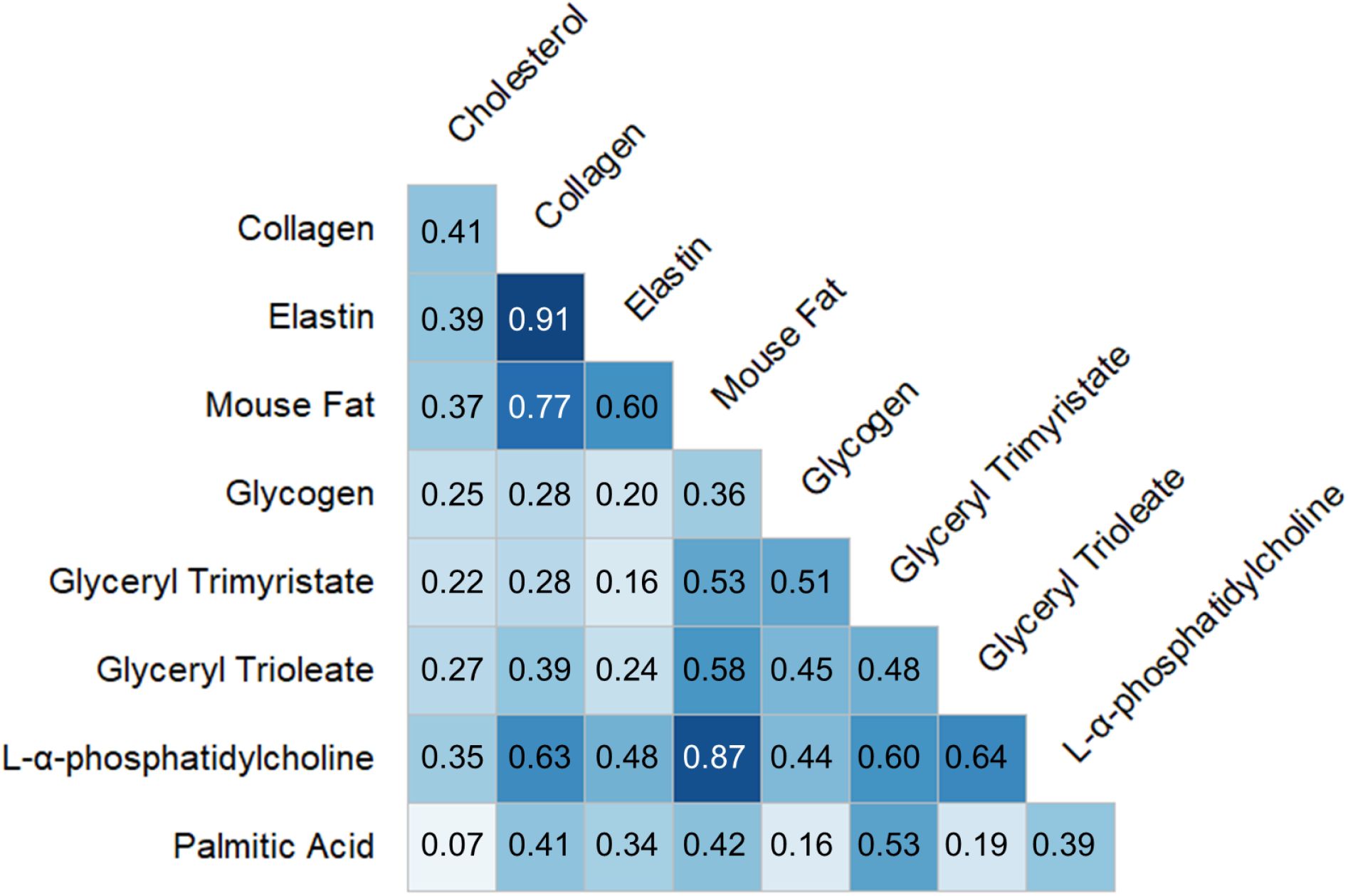
Correlation plot of pure component spectra. Due to the understanding that collagen makes up most of the uterine ECM when compared to elastin, the two proteins were combined for analysis. The excised mouse fat displayed the highest correlation with the pure l-α-phosphatidylcholine.

**Supplemental Figure 4.**
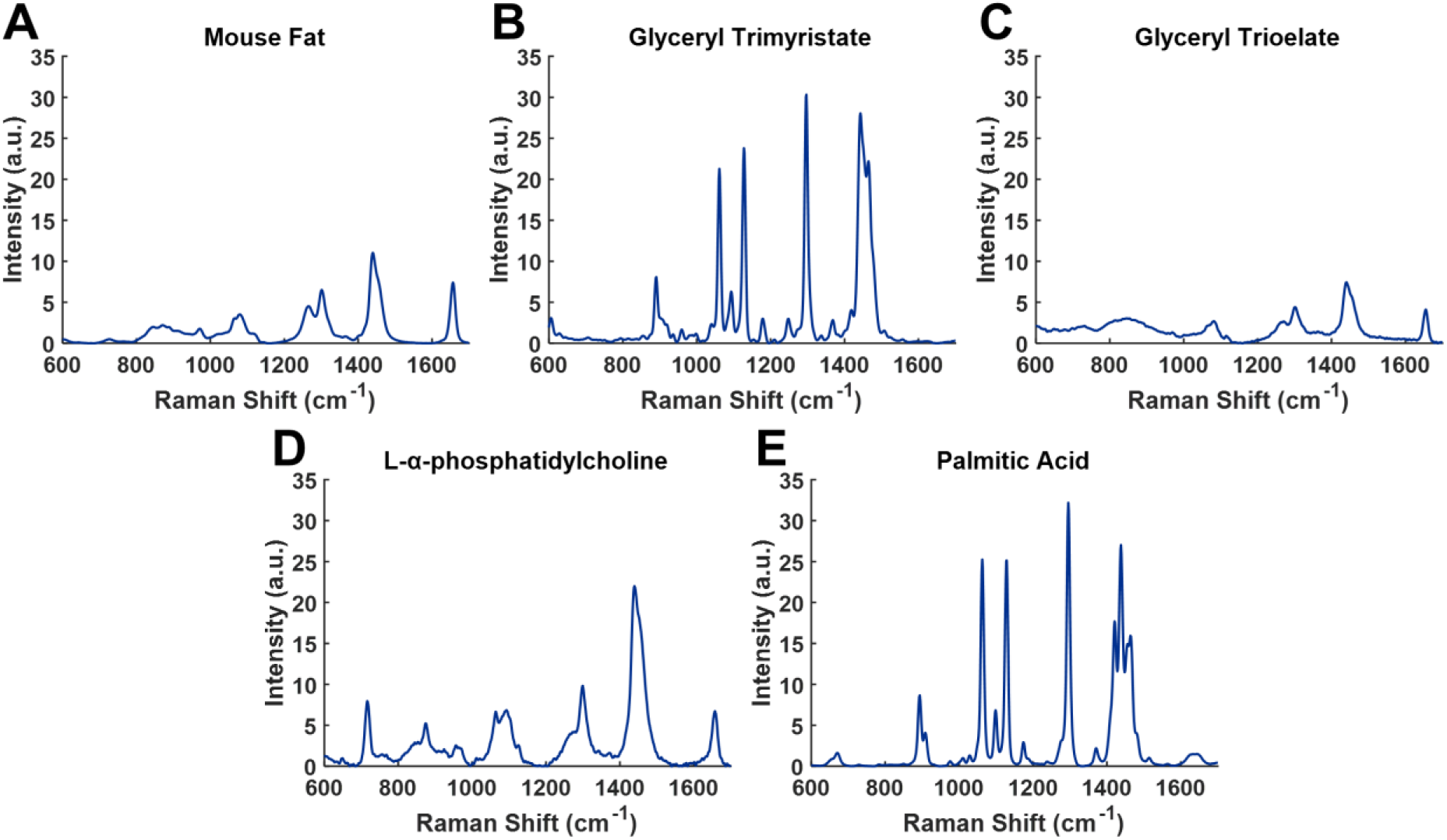
Potential spectra of fats for the NNLS model. **(A)** The fat excised from a 4-6 month murine abdominal cavity. **(B)** Pure glyceryl trimyristate (saturated triglyceride). **(C)** Pure glyceryl trioleate (unsaturated triglyceride). **(D)** Pure l-α-phosphatidylcholine (phospholipid). **(E)** Pure palmitic acid (saturated fatty acid).

**Supplemental Figure 5.**
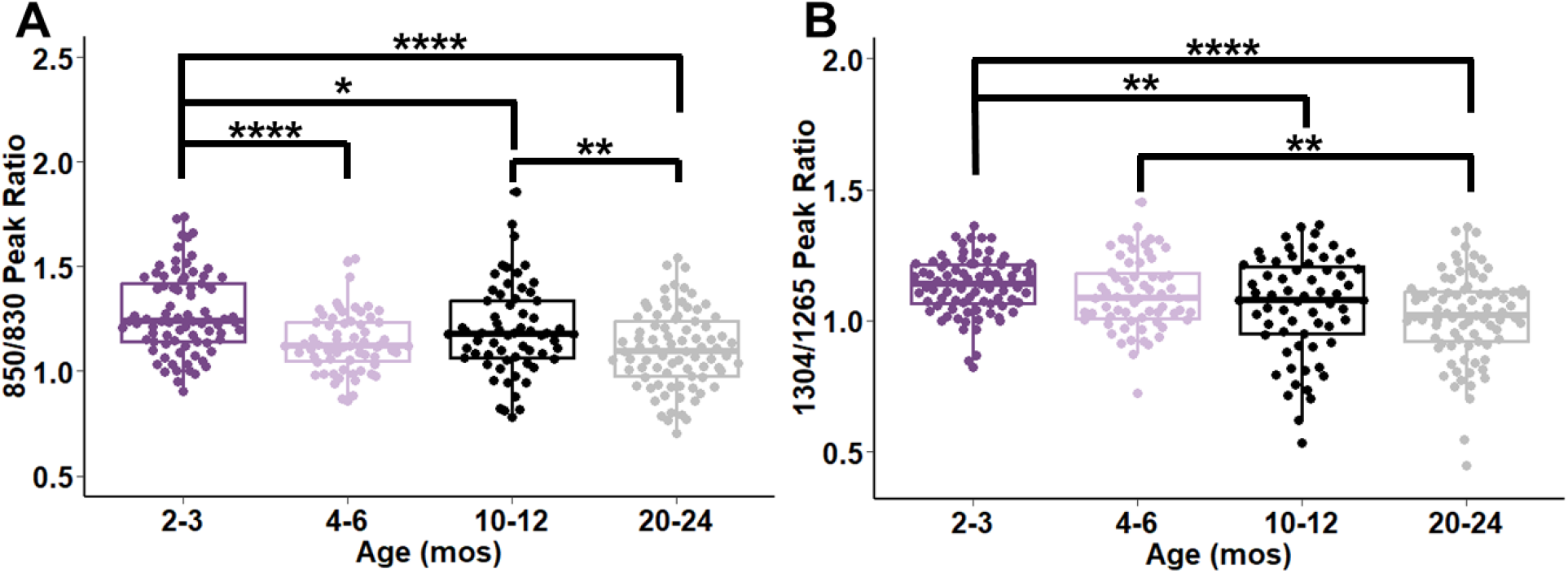
Comparison of optical peak ratios with age. **(A)** 850/830 tyrosine and **(B)** 1304/1265 peak ratios for the 2-3 month (dark purple), 4-6 month (light purple), 10-12 month (black) and 20-24 month (grey) uterus. The youngest age group showed the largest tyrosine ratio when compared to other uterine spectra (4-6 month, p<0. 0001, 10-12 month, p=0.018, 20-24 month, p<0.0001). The 20-24 month uterus displayed a significant decrease compared to the 10-12 month group (p=0.003). The 850/830 tyrosine doublet increases with cataracts (buildup of proteins in the lens of the eye) in rabbits. The youngest age group showed the highest 1304/1265 peak ratio when compared to the 10-12 month (p=0.003) and 20-24 month (p<0.0001) uterus. The oldest group displayed a significantly lower ratio compared to the 4-6 month uterus (p=0.003). The 1304 cm^-1^ peak is representative of lipid content, while the 1265 cm^-1^ peak is related to protein content [42]. ^****^p<0.0001, ^***^p<0.001, ^**^p<0.01, ^*^p<0.05 denotes statistical significance, n=8/group.

**Supplemental Figure 6.**
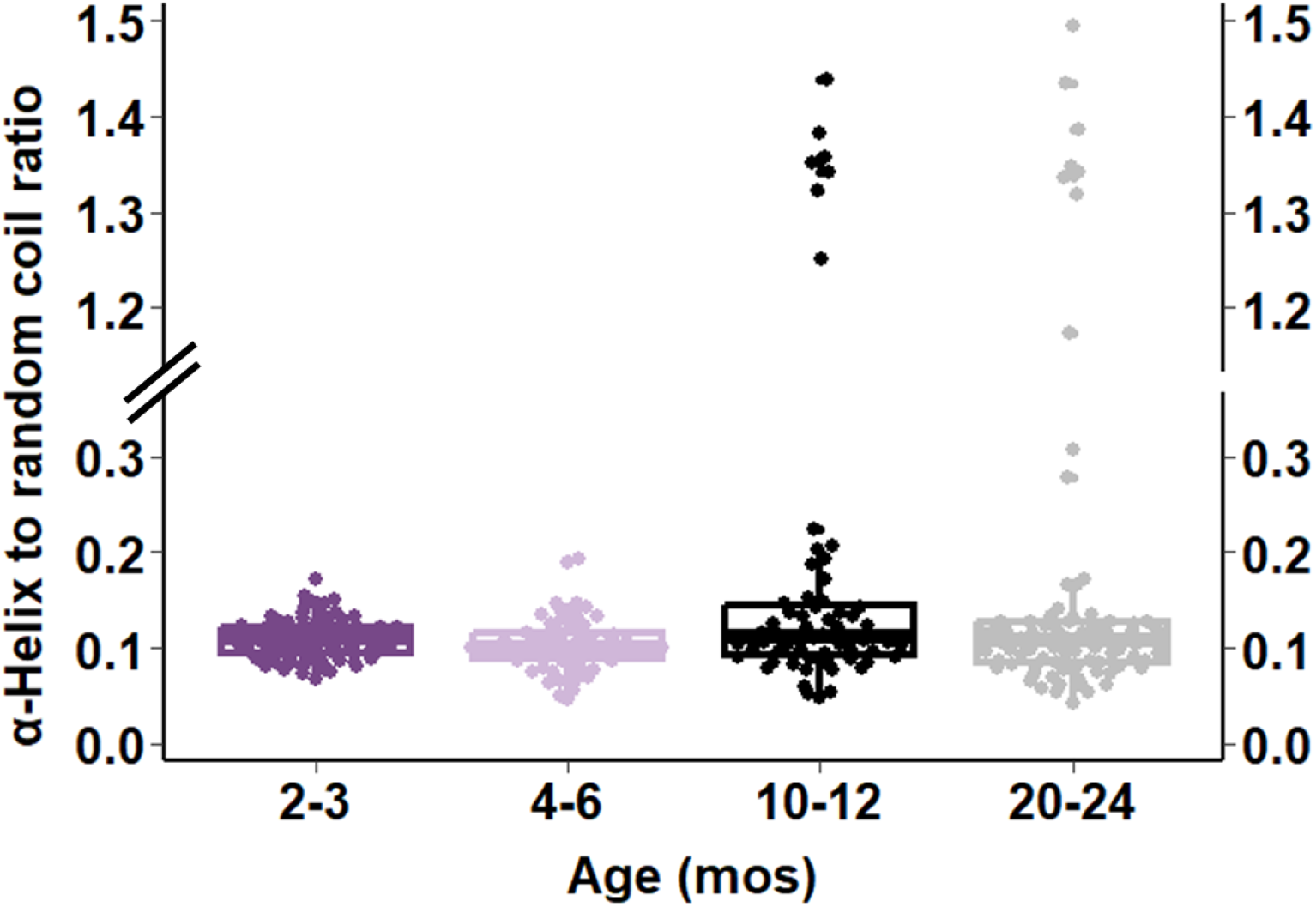
α-helix to random coil ratio for the 2-3 month (dark purple), 4-6 month (light purple), 10-12 month (black) and 20-24 month (grey) uterus. There were no significant differences between the age groups.

**Supplemental Figure 7.**
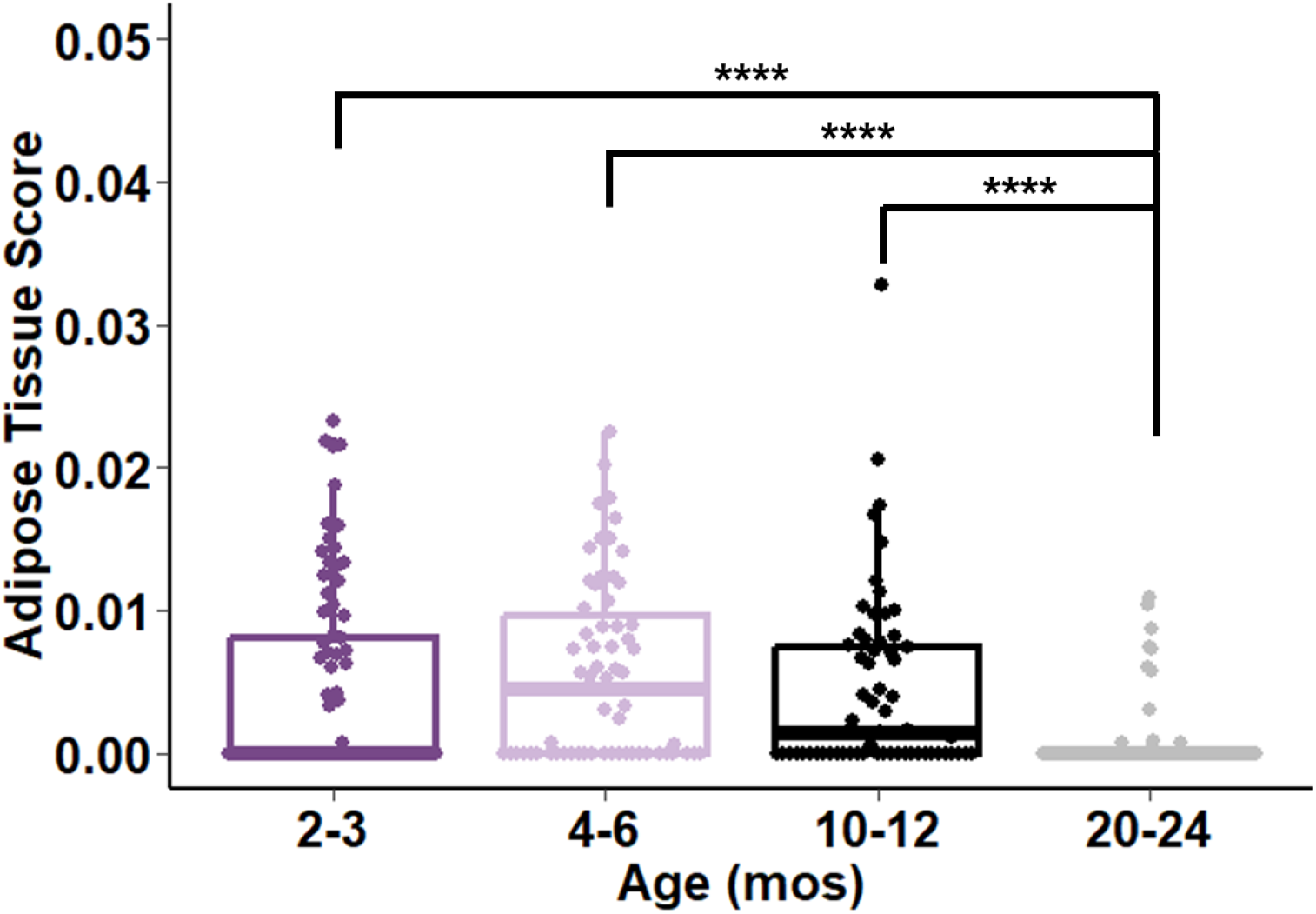
Adipose tissue NNLS score for the 2-3 month (dark purple), 4-6 month (light purple), 10-12 month (black) and 20-24 month (grey) uterus. The 20-24 month uterus displayed a significant decrease in the fat content when compared to the other age groups (p<0.0001). ^****^p<0.0001, ^***^p<0.001, ^**^p<0.01, ^*^p<0.05 denotes statistical significance, n=8/group.

## Notes

### Competing Interest Statement

The authors have declared no competing interest.

